# Feeding behavior of Costa Rican velvet worms: food hiding, parental feeding investment and ontogenetic diet shift (Onychophora: Peripatidae)

**DOI:** 10.1101/452706

**Authors:** José Pablo Barquero-González, Álvaro Vega-Hidalgo, Julián Monge-Nájera

## Abstract

We report, for the first time in onychophorans, food hiding, parental feeding investment and an ontogenetic diet shift two weeks after birth: from the parent’s adhesive used to capture prey, to the prey itself.

## Resumen

Informamos, por primera vez en onicóforos, de ocultamiento de alimentos, inversión parental alimentaria y cambio ontogénico de alimento. En las primeras dos semanas de vida, los jóvenes se alimentan exclusivamente de la goma usada por los progenitores para capturar las presas, y luego cambian a alimentarse de las presas en sí.

## Palabras clave

comportamiento alimentario, Peripatidae, comportamiento de Panarthropoda, onicóforos costarricenses no descritos, inversión parental.

The basic feeding behavior of onychophorans has been known for over a century: they capture prey with aid of an adhesive ejected by oral papillae, bite with their tooth into the exoskeleton, and ingest the partially digested tissues (Moseley, 1874). The adhesive takes about three weeks to be replenished and is thought to be metabolically valuable, since it is reingested after hunting (Read & Hughes, 1987). In the species *Euperipatoides rowelli* Reid, 1996, prey are hunted collectively and consumed in hierarchical order (Reinhard, & Rowell, 2005). Nevertheless, the feeding behavior has only been studied in adults and in a few of over 200 named species (Read & Hughes, 1987; Reinhard & Rowell, 2005; Mayer, et al., 2015). Here we report, for the first time in the phylum Onychophora, three behaviors: food hiding, parental feeding investment, and ontogenetic food shift.

From December 2016 to January 2018, individuals from eight morph-species of onychophorans from the sub-family Neopatida were housed in terraria (collection permit codes: SINAC-SE-CUSBSE-PI-R-133-2016 and SINAC-SE-CUSBSE-PI-R-015-2017). The full names and geographic origin are in Table 1 (see Barquero-González, et al., 2016, and Sosa-Bartuano, et al., 2018, for further details). For terraria we used plastic containers: 45 × 29 × 39 cm for the Gandoca species and 33 × 20 × 12 cm for the other species. Each terrarium had dirt, mosses, twigs and bark; and 1-3 females with their newborns (rarely, also one male). Settings: natural light; 99% constant humidity; 25-27°C at day and 22-23°C at night (Inkbird Thermometer & Hygrometer ITH-10).

**Table 1.** Details on individuals used for this report, and number of prey according to which prey part was consumed first. New morpho-species are indicated with *.

| **Species** | **Time kept** | **Adult individuals kept** | **Head first feeding** | **Thorax first feeding** | **Abdomen first feeding** |
| --- | --- | --- | --- | --- | --- |
| Agujas Purple Brown Onychophoran | 5 months | 2 (plus 1 offspring) | 2 | 2 | 6 |
| Batán Burgundy Brown Onychophoran | 1 month | 3 (plus 1 offspring) | 3 | 0 | 0 |
| *Epiperipatus biolleyi* | 1 month | 5 (plus 7 offspring) | 4 | 0 | 0 |
| Fortuna Burgundy Brown Onychophoran* | 2 months | 2 (plus 1 offspring) | 4 | 2 | 0 |
| Gandoca Blue Onychophoran | 5 months | 1 (plus 7 offspring) | 3 | 5 | 0 |
| *Peripatus solorzanoi* | 5 months | 3 (plus 5 offspring) | 6 | 2 | 0 |
| Quesada Burgundy Brown Onychophoran | 5 months | 3 (plus 2 offspring) | 8 | 1 | 0 |
| San Vito Collared Onychophoran* | 2 months | 2 (plus 3 offspring) | 1 | 0 | 4 |
| Sarapiquí Yellow Brown Onychophoran | 1 month | 2 (plus 1 offspring) | 2 | 0 | 0 |

They were fed once a week with live or freshly killed domestic crickets introduced from 19:00 to 20:00 h. Prey not consumed after 12 hours was removed to prevent contamination.

The onychophorans hid food in burrows, and under moss or bark: small prey was dragged, while pieces of larger prey were carried in the mouth (Fig. 1), behavior seen 3 times in the San Carlos species, 5 in Drake and 2 in Fortuna. Hiding unfinished food protects it from predators and scavengers, and has evolved independently in many species, from invertebrates to mammals (Estes, 1991; Lawrence & Newton, 1995). Food hiding may be particularly important in onychophorans because they hunt only a few times per month and are slow to process food (see Read & Hughes, 1987).

**Fig. 1.**
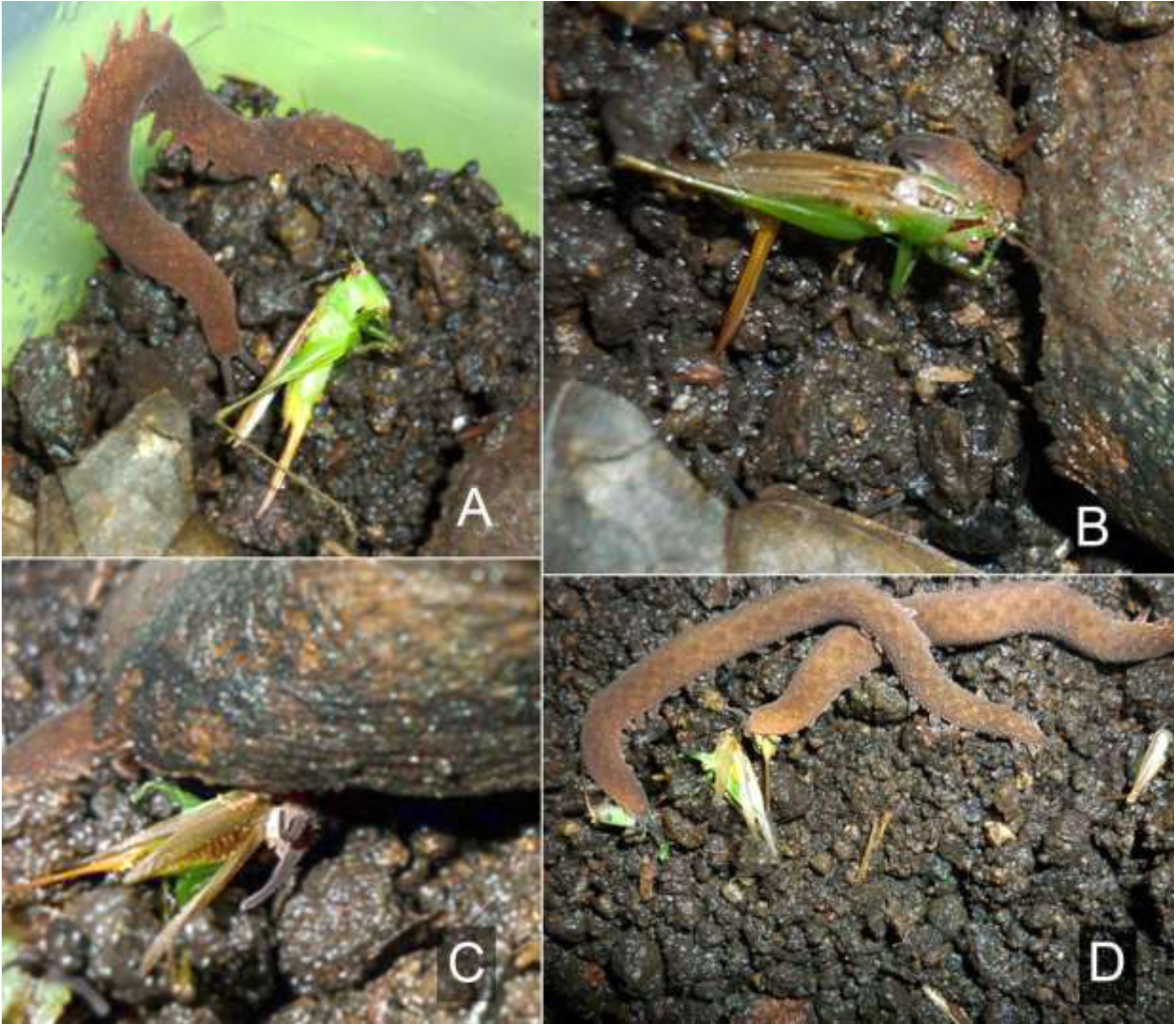
Quesada Burgundy Brown Onychophoran morph-species feeding sequence includes prey inspection (A: adult female inspecting a dead cricket that we placed in the terrarium), and hiding unfinished prey (she used her mandibles to drag it under a piece of wood, B-C). Adults can feed simultaneously, without aggression, on different prey parts (D).

Unlike *E. rowelli*, which dispute food (Reinhard & Rowell, 2005), these species shared the prey (Fig. 1D), perhaps because specimens found together tend to be relatives (see Monge-Nájera, 1995).

Specimens from Gandoca started the feeding process on the prey’s thorax (Fig. 2A); while those from Batán, Quesada, Sarapiquí and Fortuna, as well as *Epiperipatus biolleyi* and *Peripatus solorzanoi*, started feeding by the prey‘s head (Fig. 2B); the Agujas and San Vito species started by the abdominal region (Fig. 2C). The cause of this variation is ignored.

**Fig. 2.**
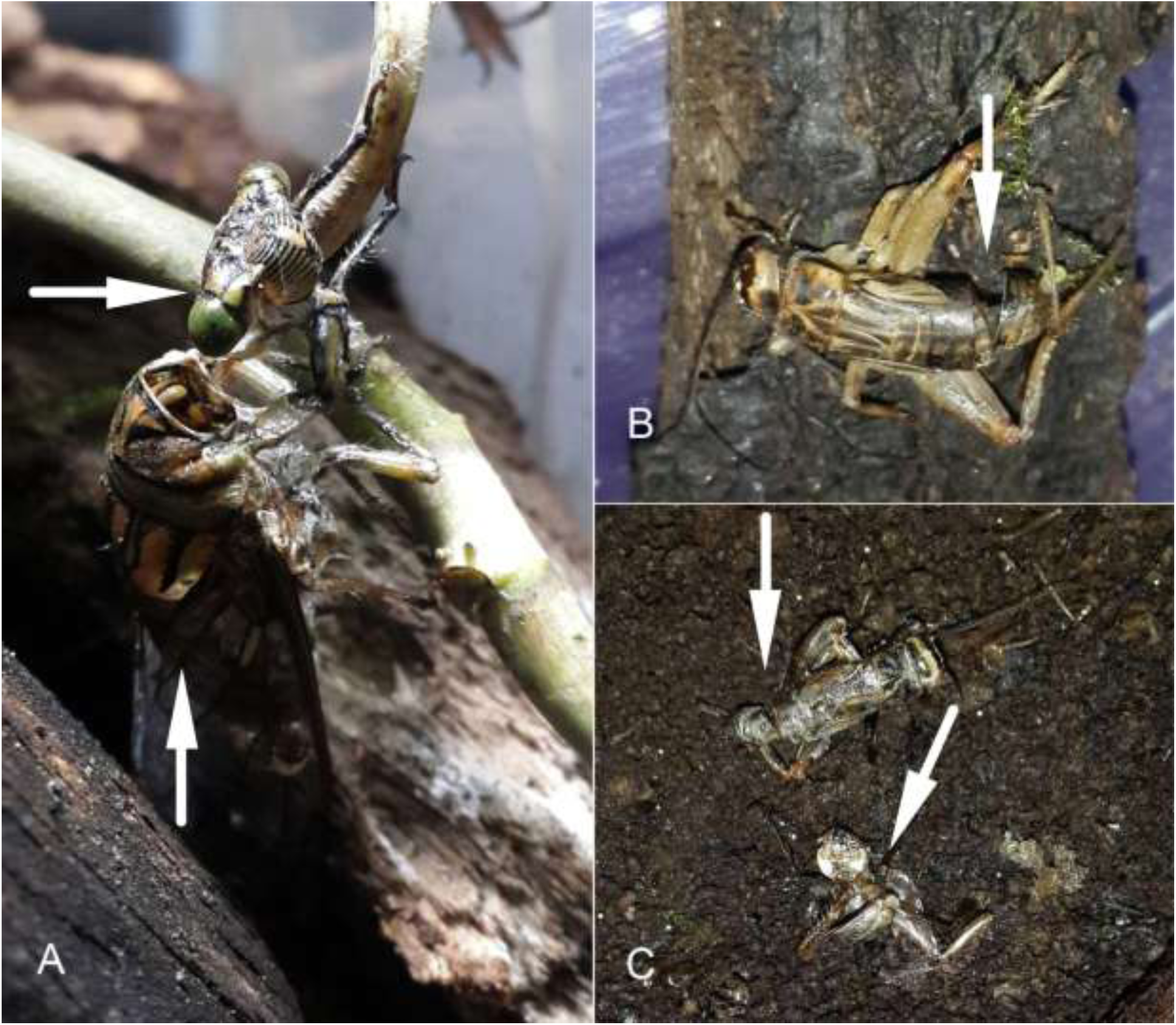
Some onychophorans start feeding by removing the head (e.g. *P. solorzanoi*, A: arrow indicates cicada head separated from thorax by the onychophoran), others by opening the abdomen (San Vito Onychophoran, B: arrow indicates gap in the abdomen), and others by separating the thorax (Fortuna Onychophoran, C: arrows indicate thorax cut in two).

During their first two weeks of life, the young only fed on the adhesive threads used to capture prey by the adults, rather than on the prey itself, apparently a form of parental feeding investment. After those two weeks, there is an ontological diet shift and adult females of the Gandocan morph-species and *P. solorzanoi*, shared the prey with their offspring (or the young found prey by themselves).

An ontological diet shift is present in a variety of organisms, and results from strong selection to optimize the use of limited resources (Brink & de Roos, 2017). To future researchers, we propose two hypotheses, that (1) the young feed on adhesive because it is a better food than the prey; or (2) that initially they cannot process prey because their digestive system is immature. Both causes are documented in other animals (Blackburn, et al., 1989; Langer, 2002) and are based on our observations of glue consumption even when prey was available to feed on.

We thank William Eberhard for advice, Bernal Morera Brenes for guidance, Diego Monge Villegas for the image editing, and three anonymous reviewers for useful comments.

